# The role of N-terminal modification of MeCP2 in the pathophysiology of Rett syndrome

**DOI:** 10.1101/122564

**Authors:** Taimoor I. Sheikh, Alexia Martínez de Paz, Shamim Akhtar, Juan Ausió, John B. Vincent

## Abstract

Methyl CpG-binding protein 2 (MeCP2), the mutated protein in Rett syndrome (RTT), is a crucial chromatin-modifying and gene-regulatory protein that has two main isoforms (MeCP2_E1 and MeCP2_ E2) due to the alternative splicing and switching between translation start codons in exons one and two. Functionally, these two isoforms appear to be virtually identical; however, evidence suggests that only MeCP2_E1 is relevant to RTT, including a single RTT missense mutation in exon 1, p.Ala2Val. Here, we show that N-terminal co- and post-translational modifications differ for MeCP2_E1, MeCP2_E1-p.Ala2Val and MeCP2_E2, which result in different protein degradation rates *in vitro*. We report partial N-methionine excision (NME) for MeCP2_E2, whereas NME for MeCP2_E1 is complete. Surprisingly, we also observed evidence of excision of multiple alanine residues from the N-terminal polyalanine stretch. Regarding MeCP2_E1-Ala2Val, we also observed only partial NME and N-acetylation (NA) of either methionine or valine. The localization of MeCP2_E1 and co-localization with chromatin appear to be unaffected by the p.Ala2Val mutation. However, a higher proteasomal degradation rate was observed for MeCP2_E1-Ala2Val compared with that for wild type (WT) MeCP2_E1. Thus, the etiopathology of p.Ala2Val is likely due to a reduced bio-availability of MeCP2 because of the faster degradation rate of the unmodified defective protein. MeCP2_E1 is thought to have a much higher translational efficiency than MeCP2_E2. Our data suggest that this increased efficiency may be balanced by a higher degradation rate. The higher turnover rate of the MeCP2_E1 protein suggests that it may play a more dynamic role in cells than MeCP2_E2.

**Significance statement:** The Rett syndrome protein, MeCP2, undergoes a number of modifications before becoming functionally active in the body’s cells. Here, we report the presence of N-terminal modifications in both MeCP2 isoforms, MeCP2_E1 and MeCP2_E2, and that the only reported Rett missense mutation in exon 1, p.Ala2Val, disrupts these modifications, decreasing the longevity of the protein. Interestingly, p.Ala2Val mutations have been reported in many other disease genes, such as *DKCX, ECHS1, IRF6, SMN1*, and *TNNI3*, and the etiopathological mechanism(s) have never been explained. Thus, this work is important not only for the understanding of the pathophysiology of Rett syndrome but also for a deeper understanding of the effects of genetic mutations at the N-terminal end of genes in general.

## Introduction

Methyl-CpG-binding protein-2, MeCP2, was originally identified by its ability to bind methylated DNA (1, 2). Mutations in the *MECP2* gene were later identified to be the cause of Rett syndrome (RTT) (3). At that time, MeCP2 was thought to be encoded by only three exons; however, an upstream non-coding first exon and a putative promoter were subsequently identified (4). The translation start site for this transcript (now termed *MECP2_E2*) in the second of the four exons, and exon 1 and most of exon 2 are within the 5’ untranslated region (UTR). However, in a splice variant of *MECP2*, termed MECP*2_E1*, exon 2 is spliced out and translation is initiated from a start codon in exon 1, thus resulting in a slightly larger protein (by 12 amino acids) with a different N-terminal end (5, 6). The remainder of the protein is identical in the two isoforms, and both contain the methyl-CpG-binding domain (MBD) and transcriptional repressor domain (TRD) (7). The MeCP2_E2 isoform has been the most widely studied, and in humans, it has 486 residues and a molecular mass of ∼53 kDa. However, MeCP2_E1 is slightly larger with 498 amino acids. Studies suggest that the mRNA expression of *MECP2_E1* is higher in brain regions than that of *MECP2_ E2* (5, 6) and that the former is the predominant isoform in the brain, except for in the thalamus (8).

Overall, the N-terminal domain (NTD) of MeCP2 (Fig. 1*A*) is uncharacterized and intrinsically disordered (9) but is highly conserved across all vertebrate groups and is identical among many higher mammalian species (10). Interestingly, among the non-mammalian vertebrate groups, only the MeCP2_E1 isoform has been identified thus far, suggesting that this isoform is the evolutionary precursor (Fig S1).

**Fig. 1.**
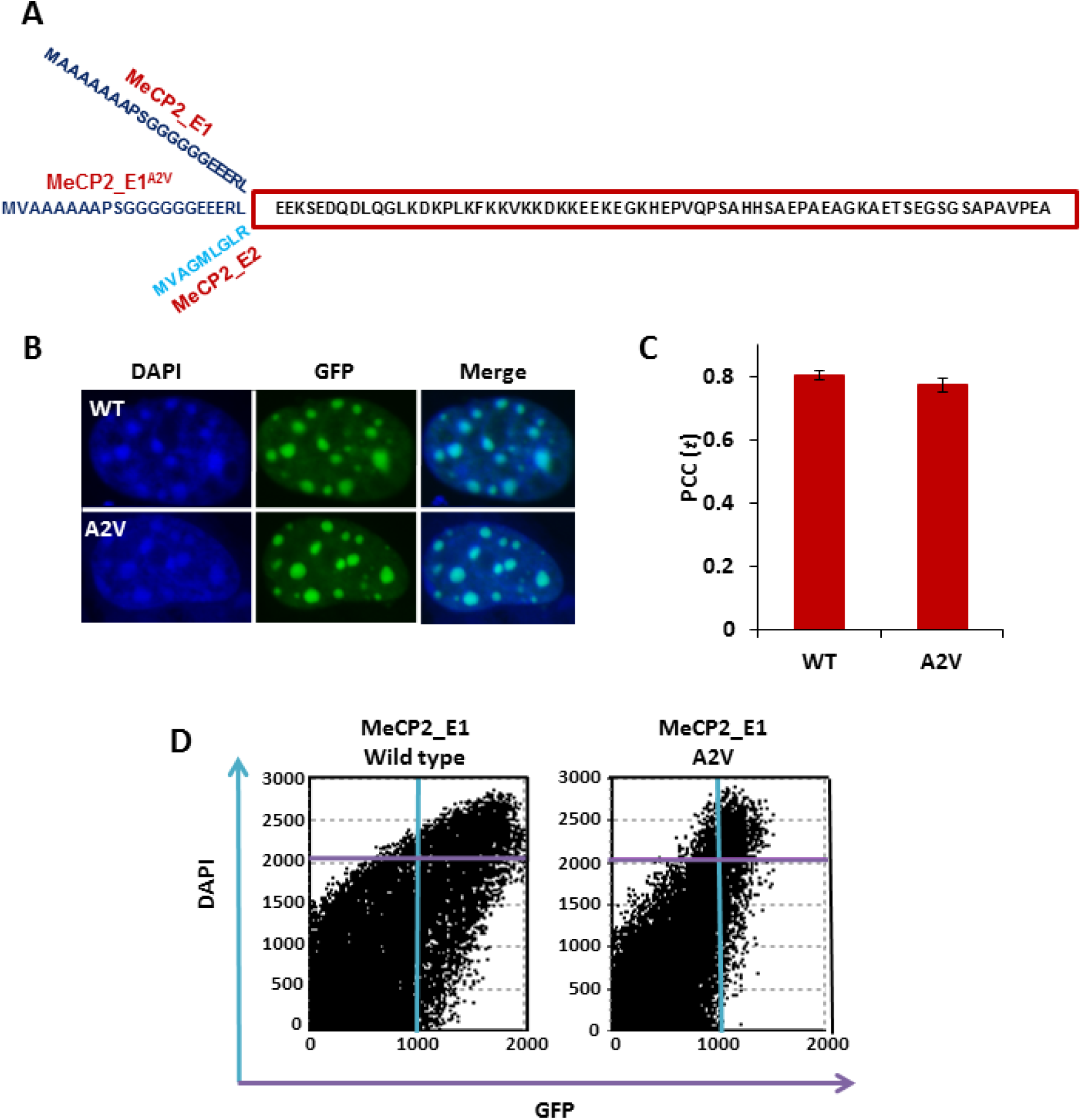
MeCP2 N-terminal domain (NTD) structure of wild type MeCP2_E1, p.Ala2Val and E2 and the effect of NTD mutation MeCP2_E1 p.Ala2Val on the translocation and localization of the protein. (A) Amino acid sequence of the MeCP2 N-Terminal Domain (NTD; residue counted as in (7)) of the mutant MeCP2_E1-p.Ala2Val and wild type MeCP2_E1/E2. (B) XY Image stacks of mouse myoblast C2C12 cells that transiently expressed the full-length MeCP2 _E1 and mutant (p.A2V), representing the co-localization at chromocenters (DAPI; column 1) and the recombinant GFP-tagged protein (column 2) and the merge of 1 and 2 (column 3). (C) Average mean of the Pearson correlation coefficient (PCC) values (n=20 cells each; ±SEM shown) of the wild type and mutant. (D) Scatter plot of blue and green pixel intensities, showing the co-localization of DNA (DAPI) and recombinant MeCP2 protein (GFP), respectively.

Studies suggest that both isoforms might have strongly overlapping functions but with spatial and temporal differences in the transcript abundance (8, 11). Complementation studies have shown that the two isoforms are able to substitute for each other and fulfill the same basic functions in the mouse brain (11). However, while the isoform-specific knockout of *Mecp2_e1* in mice recapitulates RTT-like features, the *Mecp2_e2*-specific knockout does not (12, 13). The functional role of the N-terminus of either isoform has currently not been identified. The presence of missense mutations in relation to disease is a good indicator of a functional role, and these are frequently useful for developing strategies to elucidate function. However, to date, the only exon 1 missense mutation identified in RTT patients is p.Ala2Val, which has been identified in four RTT girls and one patient diagnosed with intellectual disability (14, 15) (Rettbase entry ID # 6622 and 6622; & J. B. Vincent, *unpublished data*). This mutation would affect the E1 isoform only, leaving the E2 isoform untouched.

The removal of N-terminal methionine (NM) residues and N-terminal acetylation (NA) are the two most common protein modifications that occur either co-translationally or post-translationally (16). Methionine aminopeptidase 2 (MetAP) is the enzyme that cleaves the NM. The NM is the translational product of the start AUG codon, which is typically present only transiently on nascent protein, and its efficient removal is influenced by the penultimate residues. NM excision (NME) is much more common with smaller penultimate residues (≤ 1.29 Å radius); however, NM can be completely or partially retained if the penultimate amino acid has an intermediate-sized side chain (17). In ribosomal proteins that retain the NM, the second amino acid typically has a side chain of an intermediate size, and valine is often found at this position (18). Based on a large-scale proteomic analysis, methionine cleavage is less efficient and less likely when the second position amino acid is valine compared to when the second position is alanine (19). Overall, 100% NME has been reported in proteins with alanine at the penultimate position, whereas NME is reduced to 50% in proteins with valine at the penultimate position. An analysis of peptides with Val or Thr at the penultimate position has found NM to be less efficiently cleaved than in those with Ala, Cys, Gly, Pro, or Ser *in vitro* (19).

The addition of an acetyl group at the N-terminal residue (N-terminal acetylation; NA) usually follows directly from the removal of the NM. Among all acetylated peptides, those with a penultimate alanine are most common. However, NA cannot be definitively predicted based on the presence of Ala or Ser as terminal residues or the primary amino acid sequence (16). NA may influence the protein stability by preventing N-terminal ubiquitination (20) and may also reflect on the protein localization (21) and function. Conversely, some N-terminal acetylated proteins were found to generate a degradation signal for an ubiquitin protein known as Dao10 (22).

As the MeCP2_E1 NTD has no known ascribed function, we hypothesize that the p.Ala2Val mutation may therefore promote the retention of the NM and the reduction of NA, leading to either a reduced lifespan and/or the mislocalization of the protein. The resulting reduced bio-availability of MeCP2_E1 in girls with the p.Ala2Val mutation leads to RTT or similar phenotypes. Here, we studied the NME and NA of the two MeCP2 isoforms to understand the fundamental functional differences between them and the Ala2Val mutant MeCP2_E1 to provide a molecular link between the mutation and the disease.

## Results

### Translocation and chromatin localization of MeCP2 WT and p.Ala2Val protein

To analyze the effect of the p.Ala2Val mutation on the MeCP2-DNA interaction, we expressed C-terminal GFP-fusion protein versions of the full-length MeCP2_E1 WT and mutant in mouse C2C12 myoblast cells (chosen based on the negligible expression of endogenous MeCP2). The visual analyses of the merged images of DAPI and GFP revealed no difference in the localization of the WT and MeCP2_E1-Ala2Val protein. Pearson co-localization coefficient (PCC) values of both the MeCP2_E1 WT and MeCP2_E1-Ala2Val showed high positive correlations with *r*_*p*_ - values of 0.80 and 0.77, respectively (Fig. 1 A and B).

### *In vitro* co- / post-translational processing of WT and mutant (p.Ala2Val) MeCP2 protein

The NTDs of both MeCP2 isoforms (E1 and E2) and MeCP2_E1-Ala2Val were expressed in HEK293T cells to allow for possible N-terminal *in vitro* modifications in a mammalian cell system and were subsequently purified (Fig. S1 *A* and C). For WT MeCP2_E1, the mass spectrometry analysis found no peptides with NM, indicating complete NME at the P1 position (first residue). Moreover, acetylation of the first alanine (P’1) after NME was observed (Fig. 2*A*). Additionally, we observed several reads with the excision of the first two, three, four or six residues (*i.e*., methionine (P1) and one, two, three, or five alanines (P’1 to P’5) and acetylation of the subsequent alanine) (Fig. 2*A*). For MeCP2_E2, however, we found reads in which NM (P1) was retained and acetylated (Fig. 2*B*). For MeCP2_E2, we also found several reads with NME (P1) and acetylation of the penultimate valine (P’1) (Fig. 2*B*). Unlike WT MeCP2_E1, in sequencing MeCP2_E1-Ala2Val, we found reads with no NME (P1) but with acetylation of the N-terminal methionine (P1) (Fig. 2*C*). Additionally, a read with NME (P1) and acetylation of the penultimate valine (P’1) was also observed (Fig. 2*C*). For MeCP2_E1-Ala2Val, we also observed some evidence of NM oxidation. All PTMs reported received Ascores of 1000.

**Fig. 2.**
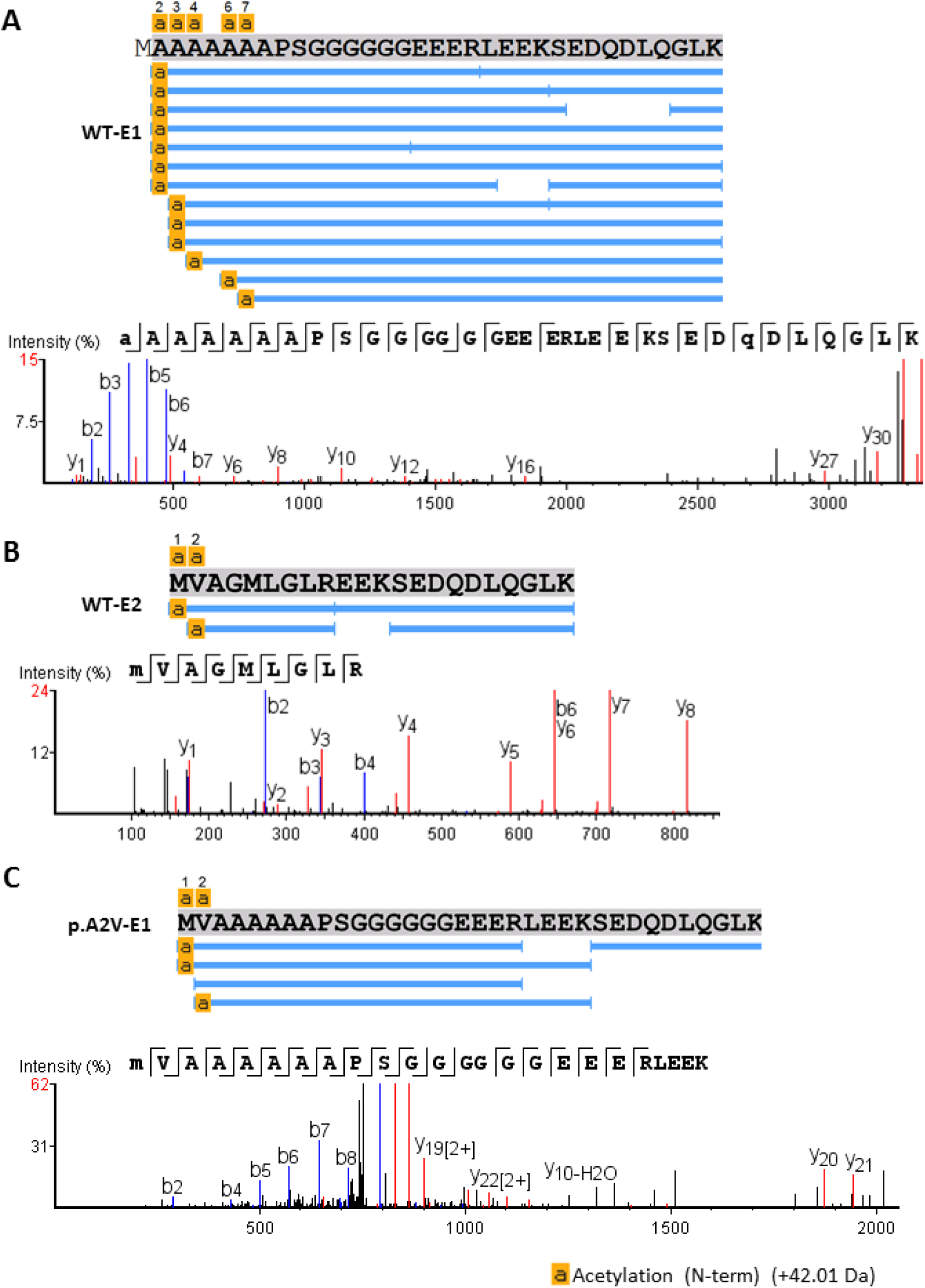
Mass spectrometry sequencing of the N-terminal of the MeCP2 protein (*in vitro*). N-terminal peptide coverage alignment chart and high-resolution mass spectra, showing N-methionine excision (NME) and N-acetylation (NA) of the N-terminus of (A) MeCP2_E1; (B) MeCP2_E2 and (C) MeCP2_E1-p.A2V. NA (+42 Da) of N-terminus amino acid is shown highlighted in yellow.

### Proteasomal degradation of MeCP2

To study the protein half-lives of MeCP2_E1, E2 and MeCP2_E1-Ala2Val, we conducted cycloheximide (CHX) chase assays. For these assays, SH-SY5Y neuroblastoma cells expressing recombinant WT MeCP2_E1, E2 and MeCP2_E1-Ala2Val GFP fusion protein were used (see Materials and Methods). MeCP2_E2 showed the longest half-life, with protein levels detected even 48 hrs after CHX treatment (Fig. 3*A*, second row). Low levels of the MeCP2_E1 protein were detected in the post-48 hrs CHX treated cells (Fig. 3*A*, first row); however, no protein was detected in the mutant cells expressing MeCP2_E1-Ala2Val 48 hrs after the CHX treatment (Fig. 3*A*, third row). The averaged band densitometric analysis of four separate experiments has been plotted (Fig. 3*B*). In similar experiments in HEK293T cells, lower levels of MeCP2_E1-Ala2Val protein were also observed compared to the wild type MeCP2_E1 and E2 protein (Fig. 3*D*, last row).

**Fig. 3.**
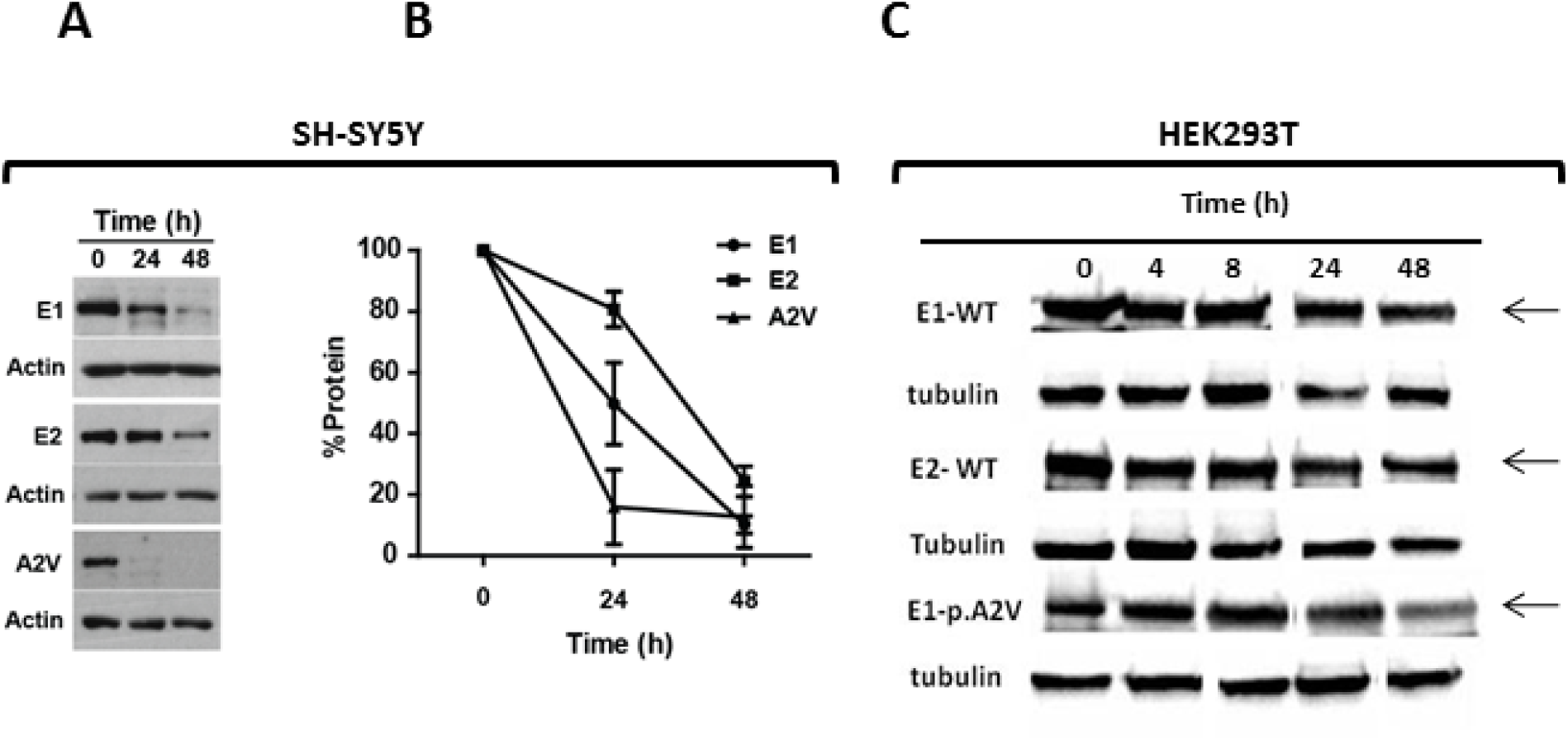
*In vitro* cycloheximide (CHX) chase assays measured the half-life of wild type MeCP2_E1 & E2 and p.A2V in cultured human cells. (A) Western blot (WB) analysis of CHX treated SK-SY5Y cells harvested after 0 (lane 1), 24 (lane 2) and 48 hrs (lane 3); (B) densitometry analysis of WBs of CHX treated SK-SY5Y cells harvested after 0, 24 and 48 hrs as shown in A; (C) WB analysis of the CHX treated HEK293T cells harvested after 0 (lane 1), 4 (lane 2), 8 hrs (lane 3), 24 hrs (lane 4) and 48 hrs (lane 5).

### Real-time mobility dynamics, degradation rates and half-life of WT and mutant (p.Ala2Val) MeCP2 protein in living cells

To further study the mobility and binding dynamics of the WT and Ala2Val mutant MeCP2 protein, fluorescence recovery after photobleaching (FRAP) assays were performed. Fig. 4*A* shows the pre-bleach, bleached and post-bleached recovery images. Surprisingly, MeCP2_E1-Ala2Val showed a slower recovery compared to WT. WT MeCP2_E1 begins to show some recovery in the 30^th^ frame, and an almost complete recovery was observed before the 100^th^ frame, whereas for Ala2Val, the recovery rate was slower and incomplete by the 100^th^ frame (Fig. 4*A*). The quantitative mean FRAP curves have shown a relatively slower fluorescent recovery of p.Ala2Val compared with that of the WT MeCP2_E1 (Fig. 4*B*). To calculate the real-time degradation rates (α) and t-half time, we performed real-time bleach-chase experiments (23, 24). HEK293T cells expressing full-length MeCP2_E1-WT and Ala2Val were used in these experiments. Bleached and unbleached cells were captured using automated time-lapse imaging for a period of 7 hrs as described in Materials and Methods. The protein degradation rates or removal rates (α) were calculated by considering the slope of the difference between the bleached, Pv(*t*), and unbleached, P(*t*) protein fluorescence on a semi-logarithmic plot using a linear regression as follows: (*ln*(*P*(*t*) − *Pv*(*t*)) (24). We observed a higher decay rate for the MeCP2_E1-Ala2Val protein compared with that of the WT (Fig. 4 C and D). For the WT, the numeric values of the protein degradation rate (α (1/h)) and t-half time were 0.014 ± 0.002 and 50.7 ± 3.5, respectively, whereas, for Ala2Val, the numeric values of the protein degradation rate (α (1/h)) and t-half time were 0.038 ± 0.002 and 18.3 ± 0.45, respectively (Fig. 4*E*).

**Fig. 4.**
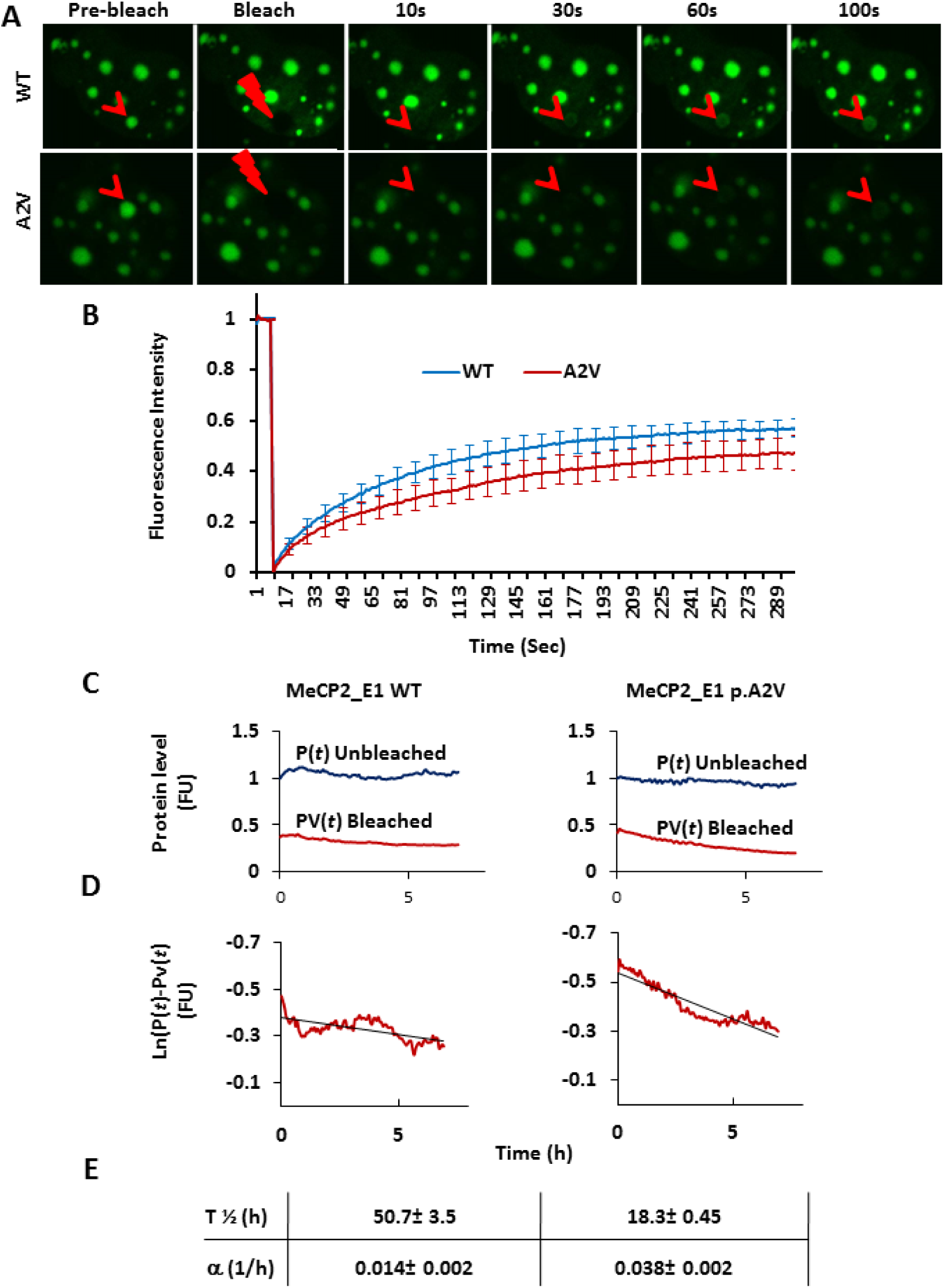
Live cell Real-Time dynamics, degradation rates and half-life of the MeCP2 protein. (A) FRAP assays, showing the recovery at different time points. (B) Fluorescent recovery of MeCP2 wild type and p.A2V. Averaged recovered intensities were plotted as a function of time from 299 image stacks (512 X 512) (SEM bar shown on every 10^th^ value; n=5). (C) Bleach chase of MeCP2_E1 and p.Ala2Val. Real-time protein recovery slope of bleached, Pv(t), and unbleached P(t) cell with the function of time (7 hrs). (D) Protein degradation rates or removal rates (α) of the same cells shown in C. (E) Protein half-life and degradation rates calculated by taking the slope of the difference between the Pv(t) and P(t) protein fluorescence on a semi-logarithmic plot using a linear regression (*ln*(*P*(*t*) − *Pv*(*t*)) (24).

## Discussion

Both human MetAPs (MetAP1 and MetAP2) preferentially cleave the N-terminal methionine of peptides, which have small penultimate residues (*i.e*., Gly, Ala, Ser, Cys, Pro, Thr, and Val), but MetAP2 has a two-fold higher cleavage activity than MetAP1 for peptides with Met-Val and Met-Thr residues at the N-terminus (25). This finding suggests that different MetAP enzymes may be required for NME of the different MeCP2 isoforms or MeCP2_E1-Ala2Val (*i.e*., MetAP2 to cleave MeCP2_E1-Ala2Val and the MeCP2_E2 isoform). In addition, reduced NME has been observed in peptides with Val or Thr at the penultimate position compared with that in peptides with Ala, Cys, Gly, Pro, or Ser at the penultimate position (19). In our results, we have observed a 100% NME of the MeCP2_E1 peptides, whereas NME for MeCP2_E2 and MeCPE1-Ala2Val was reduced, which was consistent with our predictions based on previous observations (19, 26). Additionally, for MeCP2_E1, we have also observed evidence of ‘chew back’ up to and including the first (P’1), second (P’2), third (P’3), and fifth (P’5) alanine residues of the seven-alanine polyalanine stretch after NM (P1 position). This result shows that in addition to NME, subsequent residues of MeCP2_E1 up to at least P’5 can be cleaved to generate variable N-termini. To the best of our knowledge, the cleavage of the penultimate residues in human proteins has not been reported previously. At this point, we can only speculate that such ‘chewed back’ species of MeCP2_E1 may generate subpopulations possessing different dynamics and/or stability/ longevity.

N-terminal acetylation of proteins, either with or without NME, is a common modification that involves 80-90% of human proteins (27). Several studies have described different roles of N-Ac, such as 1) regulation of cytoskeletal actomyosin interactions (28), 2) the targeting of GTPases to Golgi (29), 3) targeting to the inner nuclear membrane in yeast (30), 4) inhibition of the translocation of the protein to the endoplasmic reticulum (31), 5) acetylation of Sir3, which is required for binding to unmethylated Histone 3 Lys79, highlighting its role in gene silencing (32), 6) ubiquitination antagonist (20), and 7) as a degradation signal that is targeted by Dao10 E3 ubiquitin ligase (22) to promote degradation.

Because the NTD is just upstream of the MBD, which binds to methylated DNA (both *in vitro* and *in vivo*), to exclude the possibility that the p.Ala2Val mutation may affect nuclear translocation or chromatin localization of MeCP2, we performed co-localization experiments and calculated PCC values of MeCP2_E1-WT vs mutant MeCP2_E1-Ala2Val (CT-GFP) and DNA (DAPI) (Fig. 1 *B-D*). No significant difference in the protein-chromocenter localization was noted, indicating that mislocalization of protein is unlikely to be involved in the etiopathogenesis of p.Ala2Val. Additionally, there were no significant differences in the average number and size of chromocenters in cells transfected with the WT and mutant MeCP2_E1 constructs, which rules out an effect of the mutation on the clustering and overall chromatin organization (33).

In order to investigate the effect of the p.Ala2Val mutation on MeCP2 degradation or half-life, we conducted CHX chase experiments in which further protein translation is prevented by CHX, and MeCP2 protein degradation was monitored over time (22). In both the neuronal and non-neuronal cell types tested, both WT E1 and E2 were present even at 48 hrs post-CHX treatment. However, in neuroblastoma cells E1-Ala2Val was completely degraded by 24 hrs in the post-treatment cells (Fig. 3 *A* and C).

The bleach-chase assays, which measured the protein removal in live cells due to intracellular degradation and cell growth (23, 24), showed significantly higher removal/ degradation of MeCP2_E1-Ala2Val compared to that of the WT (Fig. 4 *C-E*), corroborating the results of the CHX assays. Since MeCP2_E1-Ala2Val shows normal localization at chromocenters (Fig. 1 *A* and *B*), the slower FRAP recovery time observed suggests that the equilibrium of chromatin-bound *versus* unbound MeCP2 has shifted as a result of its higher degradation rate. An optimum supply of correctly folded and modified MeCP2 protein is required for its role in the regulation of gene transcription, and either too little or too much MeCP2 is known to have pathogenic consequences (34, 35). Thus, disruption of the availability of MeCP2 in the nucleus through defective co-/ post-translational N-terminal modifications that lead to a faster degradation is likely to lead to dysregulation of many other essential genes, which, in turn, leads to the Rett phenotype.

Interestingly, p.Ala2Val mutations have been reported in many other disease genes, such as *DKCX* in X-linked dyskeratosis congenita (DKC1; MIM 305000) (36), *ECHS1* in mitochondrial enoyl-CoA hydratase 1 deficiency (MIM 616277) (37), *IRF6* in Van der Woude syndrome (MIM 119300) (38, 39), *SMN1* in autosomal recessive spinal muscular atrophy (MIM 253300), and *TNNI3* in familial hypertrophic cardiomyopathy type 7 (MIM 613690) (40). However, the etiopathological mechanism(s) have never been explained (36-40). Thus, the mechanistic information gained for MeCP2_E1 Ala2Val may be applicable to other diseases.

## Materials and Methods

### Cloning, mutagenesis, cell culture and transfections

C2C12 mouse myoblast cells (ATCC CRL-1772^™^) and HEK293T human embryonic kidney cells (ATCC CRL-3216™) (ATCC, Manassas, VA) were used in different experiments. Full-length and NTD mammalian expression MeCP2_E1 and MeCP2_E2 C-terminal GFP-tagged clones were prepared using vector systems, such as pcDNA3.1^™^ CT and pDEST47^™^ (Thermo Fisher Scientific, Waltham, MA). The p.Ala2Val mutation (hg19: chrX:153363118G>A; NM_001110792.1:c.5C>T) was introduced through PCR-based site-directed mutagenesis according to the manufacturer’s instructions (Quikchange Lightning site-directed mutagenesis kit, Agilent Technologies, Santa Clara, CA) using oligos ATVL-CTTCGTCCGGAAAATGGTCGCCGCC and ATVR-GGCGGCGACCATTTTCCGGACGAAG. The wild type (WT) and mutant MeCP2_E1-GFP fusion proteins were expressed in either C2C12 or HEK293T cells transfected with polyethylenimine transfection reagent (jetPEI^®^, PolyPlus, Illkirch, France) and PolyFect (Qiagen, Venlo, Netherlands), respectively, following the manufacturers’ instructions.

### Cell harvesting and protein extraction

Typically, two 150 mm plates of HEK293T cells at 80-90% confluence were used for each affinity-purification (AP) in the mass spectrometry (MS) experiments.

The cells were washed with 500 μL of ice-cold phosphate buffer saline (PBS) in each 150 mm plate and harvested with a soft sterile cell scraper. The cell pellets were washed three times by resuspending the cells gently in ice-cold PBS and centrifuging at 1500 × g at 4 °C for 5 min. The cell pellets were stored at -80 °C until use.

For the protein isolation, the pellets were resuspended in lysis buffer (50 mM HEPES-KOH, 100 mM KCl, 2 mM EDTA, 0.1% NP40, 10% glycerol, 0.25 mM Na_3_VO_4_, 50 mM β-glycerolphosphate, 1 mM NaF, and 1 mM DTT) in each plate used and incubated on ice for 30 min. The cell debris was spun for 15 min at >14-16000 × g at 4 °C. In total, 10 U DNAses I per ml of lysate (Fermentas) were added. The protein concentrations were quantified using a bicinchoninic acid (BCA) protein assay kit (Pierce^™^, Thermo Fisher Scientific, Waltham, MA). The concentration of the protein in the lysate is typically ∼5-10 mg/ml. An aliquot of lysate was also collected for the western blot analysis.

### Gel-free affinity purification / immunoprecipitation and trypsin digestion for mass spectrometry

For the gel-free GFP-purification, monovalent matrix agarose Nanobeads GFP-Trap^®^ (ChromoTek, Planegg, Germany) were used according to the manufacturer’s instruction but with the following modification (Fig. S2). Proteins were eluted 2-3 times with three bead volumes of elution buffer (0.5 M NH_4_OH, pH 11.0-12.0 (Sigma), 0.5 mM EDTA) at 4 °C with end-over-end agitation for 15 mins. The eluates were lyophilized in a centrifugal evaporator. The purified proteins were run on a 12% sodium dodecyl sulfate polyacrylamide gel electrophoresis (SDS-PAGE) gel, followed by western blotting using an anti-GFP antibody (Thermo Fisher Scientific,, Waltham, MA) to analyze the purity and assess the molecular size (Fig. S1 *A*-*B*). Prior to the mass spectrometry, the protein samples were trypsin-digested by resuspending the protein in 50 mM NH_4_HCO_3_ (pH = 8.3) and 1/10^th^ volume of a solution of 45 mM DTT. The samples were incubated at 60 °C for 30 min and allowed to cool to room temperature (RT), and then, 1/ 10^th^ of the volume of 100 mM iodoacetamide was added and incubated at RT in the dark for 15 min. The samples were then digested overnight at 37 °C with 0.75 μg of porcine trypsin, and an additional 0.75 μg of trypsin was added the next day to continue the digestion for another 3 hours. The samples were lyophilized and stored at -80 °C until use in the mass spectrometry experiment.

### Mass spectrometry to determine the MeCP2 PTMs

The protein samples were digested overnight at 37 °C with trypsin at a 50:1 protein:enzyme ratio. After desalting the peptide mixtures using C_18_ reverse phase columns, the tryptic peptides were loaded onto a 50 cm × 75 μm ID column with RSLC 2 μm C_18_ packing material (EASY-Spray, Thermo-Fisher, Odense, Denmark) with an integrated emitter. The peptides were eluted into a Q-Exactive™ Hybrid Quadrupole-Orbitrap^™^ mass spectrometer (Thermo Fisher Scientific, Waltham, MA) using an Easy-Spray nLC 1000 chromatography system (Thermo Fisher Scientific, Waltham, MA) with a 1 hr gradient from 0% to 35% acetonitrile in 0.1% formic acid. The mass spectrometer was operated in a data-dependent mode with 1 mass spectrometry (MS) spectrum, followed by 10 MS/MS spectra. The MS was acquired with a resolution of 70,000 FWHM (full width at half maximum), a target of 1 x 10^6^ ions and a maximum scan time of 120 ms. The MS/MS scans were acquired with a resolution of 17,500 FWHM, a target of 1 X 10^6^ ions and a maximum scan time of 120 ms using a relative collision energy of 27%. A dynamic exclusion time of 15 seconds was used for the MS/MS scans. The raw data files were acquired with an Xcalibur 2.2 (Thermo Fisher Scientific, Waltham, MA) and processed with the PEAKS 7 search engine (Bioinformatics Solutions, Waterloo, ON) using a database consisting of the wild type and mutant constructs of MeCP2. The identified peptides and PTMs were assigned Ascores by the PEAKS software (39).

### Cell fixation and Confocal Imaging

For the fixed cell imaging, the cells were fixed using 4% paraformaldehyde, followed by two washes with ice-cold phosphate buffer saline (PBS), and the transfected cells were then counter stained with DAPI (4′-6′-diamidino-2-phenylindol). Confocal microscopy was used to study the colocalization of MeCP2 accumulation at chromocenters. Z-stacks were acquired with a frame size of 512 × 512 pixels using the GFP^488^ and DAPI^405^ channels.

### Single cell nuclei identification

The identification of the nuclei of single cells and chromocenters was performed by intensity-based thresholding and the implementation of the Water algorithm (41). Chromocenters and GFP-fusion protein co-localization images were captured by confocal microscopy (Olympus FV-1200) using a 63×/1.3 NA oil objective at 405 nm diode pumped solid state for DAPI, and 488 nm argon for GFP.

### Fluorescence recovery after photobleaching

To study the protein recovery and dynamics in living cells, MeCP2_E1 WT and mutant (p.Ala2Val) were expressed in transfected C2C12 cells in chambered cover glass culture plates (Nunc^™^; NalgeNunc, Rochester, NY). The fluorescence recovery after photobleaching (FRAP) assay was used to capture the recovery and diffusion dynamics. Five independent FRAP experiments in temperature (37 °C) and CO_2_ (5%) controlled incubation chambers were performed on the WT and mutant recombinant proteins in the C2C12 cells.

For the time-lapse imaging, a series of confocal time-lapse images of frames (512 × 512 pixels) imaged at 488 nm laser excitation with 0.05 transmission were used to record the GFP-tagged protein post-bleach recovery. The FRAP assays were recorded with a minimum of 25 pre-bleach frames, 200 μs bleach time with 405 nm laser line at 100% transmission, and 299 post-bleach frames were recorded at equal time intervals.

### Cycloheximide chase assay

SH-SY5Y neuroblastoma cells were transfected with MeCP2_E1, MeCP2_E2 and MeCP2_E1-p.Ala2Val cloned in the GFP fusion expression vectors as described above. The cells were seeded one day before transfection in the culture dishes. After 24 hrs, the cells were transfected using Lipofectamine 3000. At 48 hrs post-transfection, the cells were treated with 10 μg/mL of cycloheximide (CHX) and harvested after 0, 24 and 48 hrs. The cell lysates were prepared using 10 mM Tris/Cl pH 7.5; 150 mM NaCl; 0.5 mM EDTA; 0.5% NP-40, 0.02% Thimerosal (preservative) lysis buffer. Denaturing SDS-PAGE followed by western blot analysis were used to detect the GFP-fused recombinant protein using an anti-GFP tag antibody (Thermo Fisher Scientific, Waltham, MA). All experiments were conducted in four replicates to confirm the reproducibility.

### Real-time bleach-chase assays

To study the protein degradation / removal rate and real-time half-life in living cells, MeCP2_E1 WT and mutant (p.Ala2Val) were expressed and transfected into HEK293T cells in 35 mm dishes with No. 1.5 coverslips (MatTek Corporation MA USA). All bleach-chase experiments were conducted in temperature (37 °C)- and CO_2_ (5%)-controlled incubation chambers.

A series of confocal time-lapse images at 20x resolution (frame size of 512 × 512 pixels) were captured at 488 nm laser excitation with 0.05 transmission. Half of the cells in the field of objective were bleached using a 405-nm laser for 250 frames (200 μs bleach time per frame). The time-lapse images of the visible fluorescent protein after bleaching (bleached cell, Pv(*t*)) and the total fluorescent protein (unbleached cells, P(*t*)) were recorded for ∼7 hrs (a total of 105 frames) with ∼4-min intervals.

## Data analysis

The Olympus FluoView (FV1200) software was used to captured the cell images and calculate the Pearson’s correlation coefficient (PCC) for the co-localization of the chromocenters (DAPI) and recombinant protein (GFP). A MATLAB-based Windows application, easyFRAP (42), was used to analyze the FRAP data. The descriptive statistical data analysis was calculated using Microsoft Excel and two-tailed unpaired Student’s *t*-test, and the *p*-values for statistical significance were calculated using GraphPad^™^ software online tools.

## Acknowledgments

This work was supported by a grant from the Ontario Rett Syndrome Association (ORSA) to JV and JA, by donations made to the Centre for Addiction and Mental Health Foundation, and a Canadian Institutes of Health Research (CIHR) grant (MOP-130417]) to JA. TIS was supported by awards from the Dalton Whitebread Scholarship Fund and from the Margaret and Howard Gamble Research Grant.

